# The impact of dominance on adaptation in changing environments

**DOI:** 10.1101/2020.07.13.200642

**Authors:** Archana Devi, Kavita Jain

## Abstract

Natural environments are seldom static and therefore it is important to ask how a population adapts in a changing environment. We consider a finite, diploid population evolving in a periodically changing environment and study how the fixation probability of a rare mutant depends on its dominance coefficient and the rate of environmental change. We find that in slowly changing environments, the effect of dominance is the same as in the static environment, that is, if a mutant is beneficial (deleterious) when it appears, it is more (less) likely to fix if it is dominant. But in fast changing environments, the effect of dominance can be different from that in the static environment and is determined by the mutant’s fitness at the time of appearance as well as that in the time-averaged environment. We find that in a rapidly varying environment which is neutral on average, an initially beneficial (deleterious) mutant that arises while selection is decreasing (increasing) has a fixation probability lower (higher) than that for a neutral mutant as a result of which the recessive (dominant) mutant is favored. If the environment is beneficial (deleterious) on average but the mutant is deleterious (beneficial) when it appears in the population, the dominant (recessive) mutant is favored in a fast changing environment. We also find that when recurrent mutations occur, dominance does not have a strong influence on evolutionary dynamics.

## 1 Introduction

Natural environments change with time and a population must continually adapt to keep up with the varying environment (Gillespie, 1991; Messer *et al.*, 2016; Bleuven and Landry, 2016). It is therefore important to understand the adaptation dynamics of a finite population subject to both random genetic drift and environmental fluctuations. This is, in general, a hard problem but some understanding of such dynamics has been obtained in previous investigations. For example, when the environment changes very rapidly, on the time scale of a generation, the dynamics of adaptation are simply determined by the time-averaged environment (Gillespie, 1991).

Environments can, of course, vary slowly and recent experiments have shown the impact of rate of change in the environment on the population fitness (Salignon *et al.*, 2018; Boyer and Sherlock, 2019). The fixation probability, fixation time and adaptation rate in changing environments have also been studied in a number of theoretical studies (Takahata *et al.*, 1975; Gillespie, 1993; Mustonen and LÄssig, 2008; Assaf *et al.*, 2008; Uecker and Hermisson, 2011; Waxman, 2011; Peischl and Kirkpatrick, 2012; CvijoviĆ *et al.*, 2015; Dean *et al.*, 2017), and it has been found that when the environment changes at a finite rate, the population dynamics are strongly determined by the environment in which the mutant arose.

In diploid populations, the degree of dominance also affects the evolutionary dynamics; in particular, in a static environment, the fixation probability of a dominant beneficial mutant is known to be higher than when it is recessive (*Haldane’s sieve*) (Haldane, 1927), while the opposite trend holds if the mutant is deleterious (Kimura, 1957). How these results are affected in changing environments is, however, not completely understood.

To understand the adaptive process in variable environments, Uecker and Hermisson (2011) developed a framework for general time-dependent selection schemes; however, their analysis was limited to very large populations and did not explicitly address how dominance affects the fixation process. In this article, we employ their formalism to study the dynamics of adaptation of a finite, diploid population evolving in an environment that changes periodically due to, for example, seasonal changes or drug cycling with a particular focus on the impact of dominance. We find that in time-dependent environments, the magnitude of the fixation probability of a rare mutant differs substantially from the corresponding results in the time-averaged environment. Furthermore, the dependence of the fixation probability on dominance coefficient can differ from that expected in the static environment depending on the rate of environmental change, the time of appearance of the mutant and its fitness in the time-averaged environment. However, when recurrent mutations occur, our results for the average allele frequency and population fitness suggest that dominance does not have a strong influence on evolutionary dynamics.

## 2 Model

We consider a finite, randomly mating diploid population of size *N* with a single biallelic locus under selection. The (Wrightian) fitness of the three genotypes denoted by *aa*, *aA*, *AA* is 1 + *s*, 1 + *hs*, 1, respectively, where the dominance coefficient 0 < *h* ≤ 1. The population evolves in a periodically changing environment that is modeled by a time-dependent selection coefficient 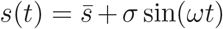 where 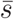 is the selection coefficient averaged over a period 2*π*/*ω*; in the following, we assume that 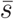 is arbitrary but *σ* > 0. We ignore random fluctuations in the environment so that selection changes in a predictable fashion. Although dominance can evolve with time (Mayo and BÜrger, 1997), for simplicity, here we assume the dominance coefficient to be constant in time. We also allow mutations to occur with a constant, symmetric probability *μ* between the two alleles.

The population dynamics are described by a continuous time birth-death model. Although in a finite population with overlapping generations, Hardy-Weinberg equilibrium (HWE) does not strictly hold, it is a good approximation when selection and mutation are weak and population size is large (Nagylaki, 1992). In the following, we therefore work in these parameter regimes and assume that the population is in HWE immediately after mating (see also Supporting Information, Sec. S1). If the birth rate of an individual is given by its genotypic fitness and the death rate by one, the number of allele *a*(*A*) increases by one if it is chosen to give birth at rate equal to its marginal fitness, *w_a_*(*w_A_*) and an allele *A*(*a*) is chosen to die. Taking the effect of mutations on these birth and death processes into account, we find that the rate *r_b_* and *r_d_* at which the number *i* of allele *a* increases or decreases, respectively, by one are given by

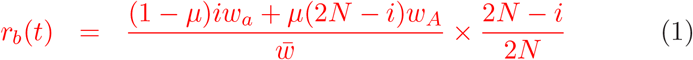

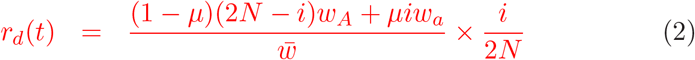

where *w_a_* = (1 + *s*)*x* + (1 + *hs*)(1 − *x*), *w_A_* = (1 − *x*) + (1 + *hs*)*x*, 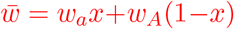 is the population-averaged fitness and *x* = *i*/(2*N*) is the frequency of allele *a*.

We study the model described above analytically using an appropriate perturbation theory and numerically through stochastic simulations in which the time interval Δ between successive generations, *t* and *t* + Δ is treated as a random variable. We first choose Δ from an exponential distribution with rate 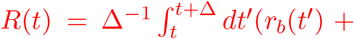 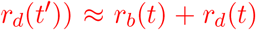 (the latter approximation is justified for small cycling frequencies as the correction to it is of order *ω*). Then the number of allele *a* and *A* is changed with a probability proportional to the rate of the respective event given above (Desai and Fisher, 2007; Uecker and Hermisson, 2011).

Below we first consider the weak mutation regime (2*Nμ* ≪ 1) in which once a mutant has appeared in a clonal population, further mutations may be ignored until the mutant is fixed or lost; here, we are interested in understanding how the fixation probability of a rare mutant depends on various environmental and population factors such as the cycling frequency and dominance parameter. We also briefly explore the strong mutation regime (2*Nμ* ≫ 1) in which recurrent mutations occur; our objective is to understand the effect of mutations and environmental fluctuations on population fitness and the dynamics of allele frequency.

## 3 Fixation probability of a rare mutant

In a static environment, the fixation probability of a mutant allele in a large population (*N* → ∞) under weak selection and weak mutation (*s, μ* → 0) with finite 2*Ns*, 2*Nμ* can be described by a backward Kolmogorov equation (Ewens, 2004). For the model described in the last section, the probability *P*_fix_ that mutant allele *a* present in frequency *x* at time *t* fixes eventually is given by (Uecker and Hermisson, 2011)

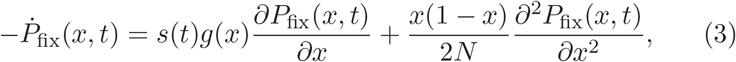

where dot denotes a derivative with respect to time and *g*(*x*) = *x*(1 − *x*)(*x*+*h*(1−2*x*)). On the right-hand side (RHS) of the above equation, the first term describes the deterministic rate of change in the allele frequency (Ewens, 2004) and the second term captures the stochastic fluctuations due to finite population size. Since the left-hand side (LHS) of (3) is nonzero, it is *inhomogeneous* in time; that is, the eventual fixation probability depends on the time of appearance *t_a_* (or phase *θ_a_* = *ωt_a_*) of the mutant (Uecker and Hermisson, 2011; Waxman, 2011).

Equation (3) does not appear to be exactly solvable, and approximate methods such as perturbation theory require an exact solution of the unperturbed problem (*σ* = 0) which is not known in a closed form for nonzero 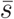 (Kimura, 1957; Jain and Devi, 2020). Therefore to obtain an analytical insight, we study the fixation probability using a branching process for positive 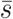 and analyze the above diffusion equation for 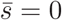 using a time-dependent perturbation theory. Some numerical results for negative 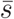 are also given. Before proceeding to a quantitative analysis, we first give a qualitative picture of the process in the following section.

### 3.1 Qualitative features

In a static environment, a dominant beneficial (deleterious) mutant has a higher (lower) chance of fixation (Kimura, 1957). In a changing environment, we expect that this result continues to hold when the mutant is beneficial or deleterious at all times (that is, when 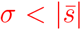 for the periodically changing *s*(*t*); see Sec. S2). But it is not obvious how dominance influences the fixation probability when the mutant is transiently beneficial or deleterious. Our results for the fixation probability in such situations are shown in Fig. 1, Fig. 2 and Fig. 3 when the mutant arises in an environment which is beneficial, neutral and deleterious on average, respectively.

**Figure 1:**
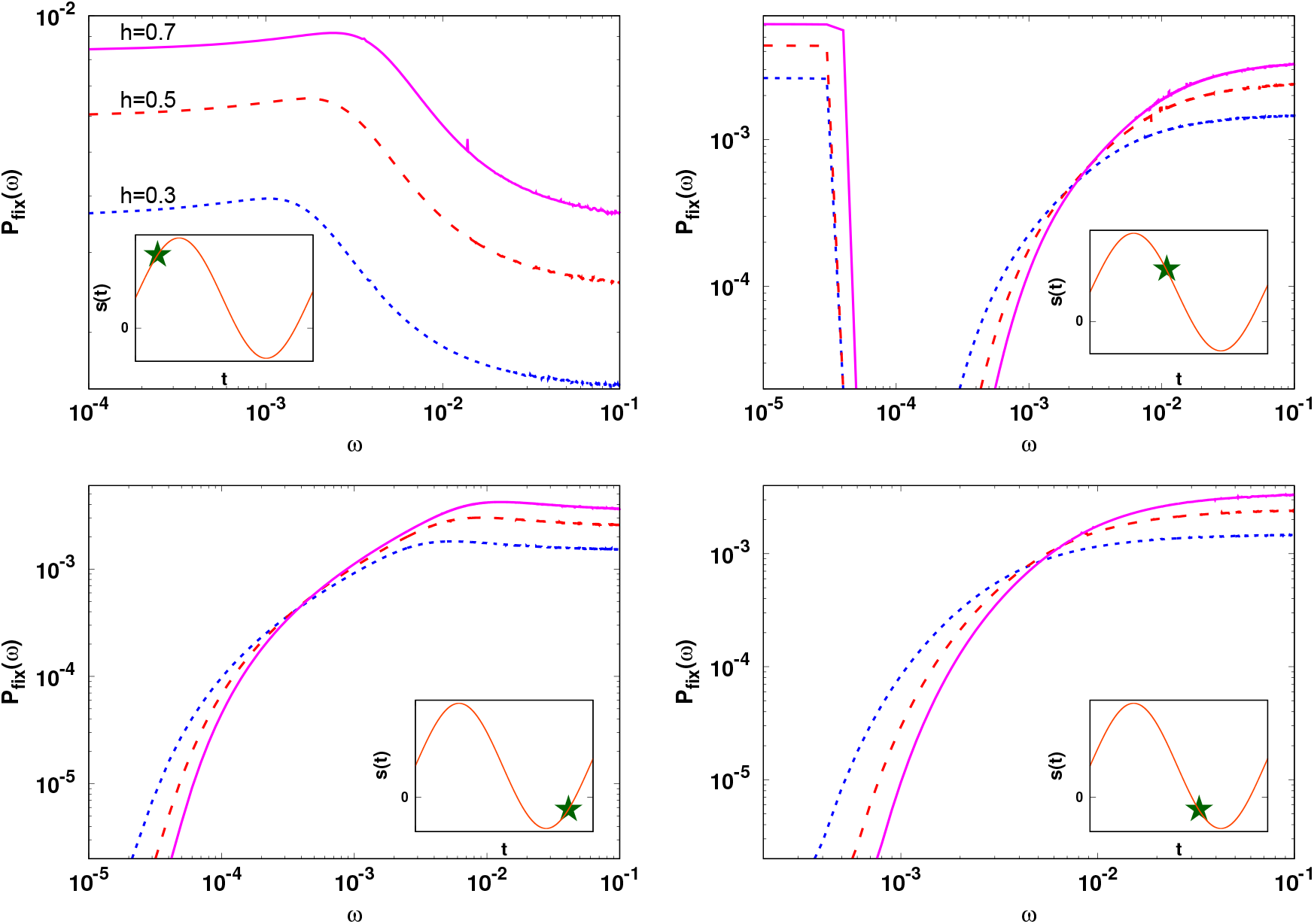
Fixation probability (5) obtained using branching process approximation for a mutant that is beneficial on average 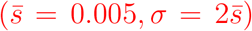 for *θ_a_* = *π*/4, 7*π*/8, 5*π*/4, 7*π*/4 (clockwise from top left as shown in insets) for *h* = 0.3 (dotted), 0.5 (dashed), 0.7 (solid). Note that the *h*-dependence of the fixation probability is different at small and large cycling frequencies when the mutant is deleterious at the time of appearance.

**Figure 2:**
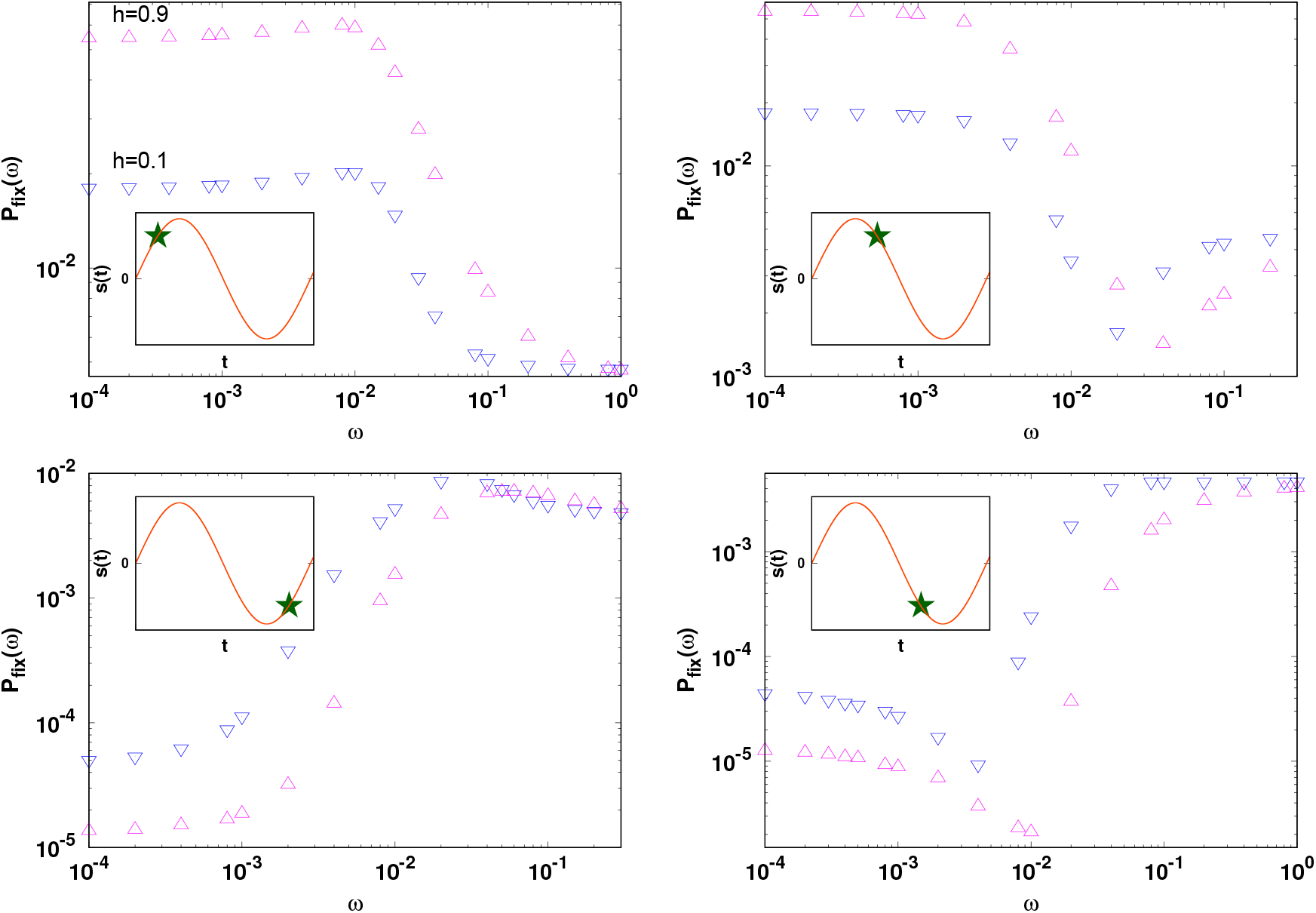
Fixation probability as a function of cycling frequency for a mutation that is neutral on average and arises in a large population of size *N* ≫ *σ*^−1^ for *h* = 0.1(▽) and 0.9(△), and *θ_a_* = *π*/4, 3*π*/4, 5*π*/4, 7*π*/4 (clockwise from top left). The other parameters are *N* = 100 and *σ* = 0.1. The simulation data are obtained by averaging over 10^7^ independent runs. A dominant (recessive) mutant that is beneficial (deleterious) when it appears in the population can be disfavored at large cycling frequencies.

**Figure 3:**
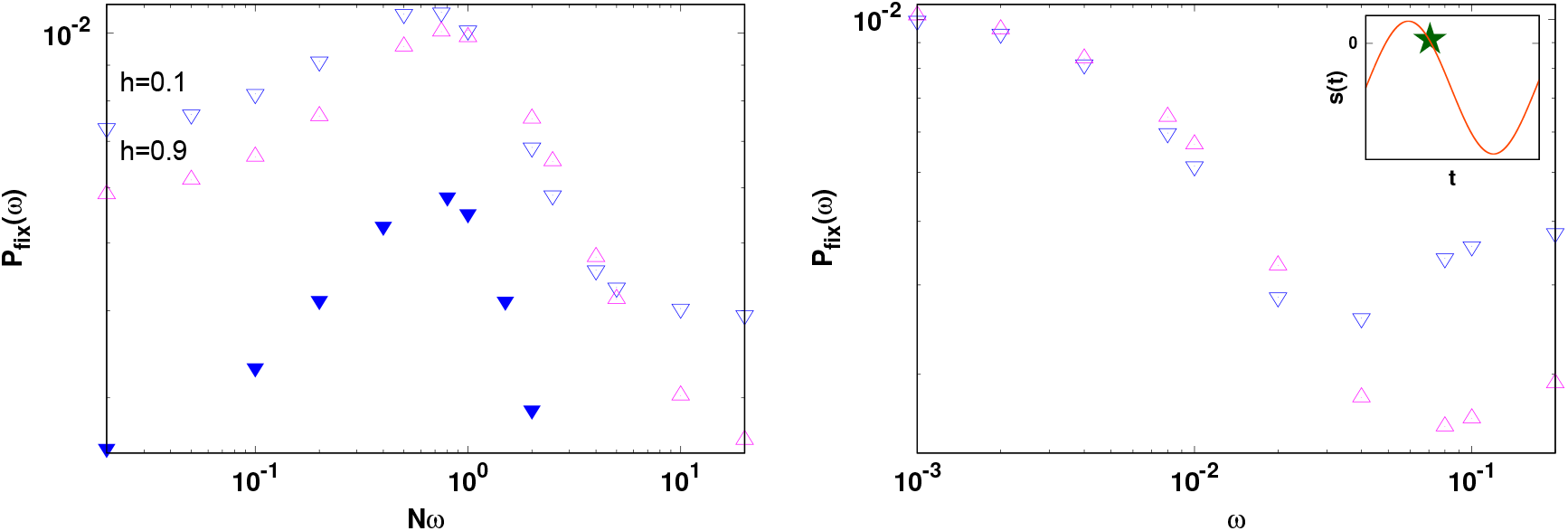
Fixation probability of a mutant which is deleterious on average for 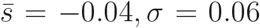, and *θ_a_* = *π*/8 (left panel) and 3*π*/4 (right panel). The simulation data are averaged over 10^7^ runs. The other parameters are *N* = 50, *h* = 0.1(▽), 0.9(△). In the left panel, the data for *h* = 0.1, *N* = 100 (▼) is shown to support the claim that the resonance frequency scales as *N* ^−1^. Note that the recessive mutant that is beneficial when it appears in the population is favored at large cycling frequencies.

In all these cases, in a slowly changing environment, the effect of dominance is found to be the same as in the static environment (that is, a dominant mutant that starts out as a beneficial (deleterious) one has a higher (lower) chance of fixation). This is because when the environment changes infinitesimally slowly (*ω* → 0), as Fig. 4a illustrates for on average neutral mutation, fixation occurs rapidly compared to the fluctuations in selection so that the sign of selection remains the same from origination to fixation of the mutant.

**Figure 4:**
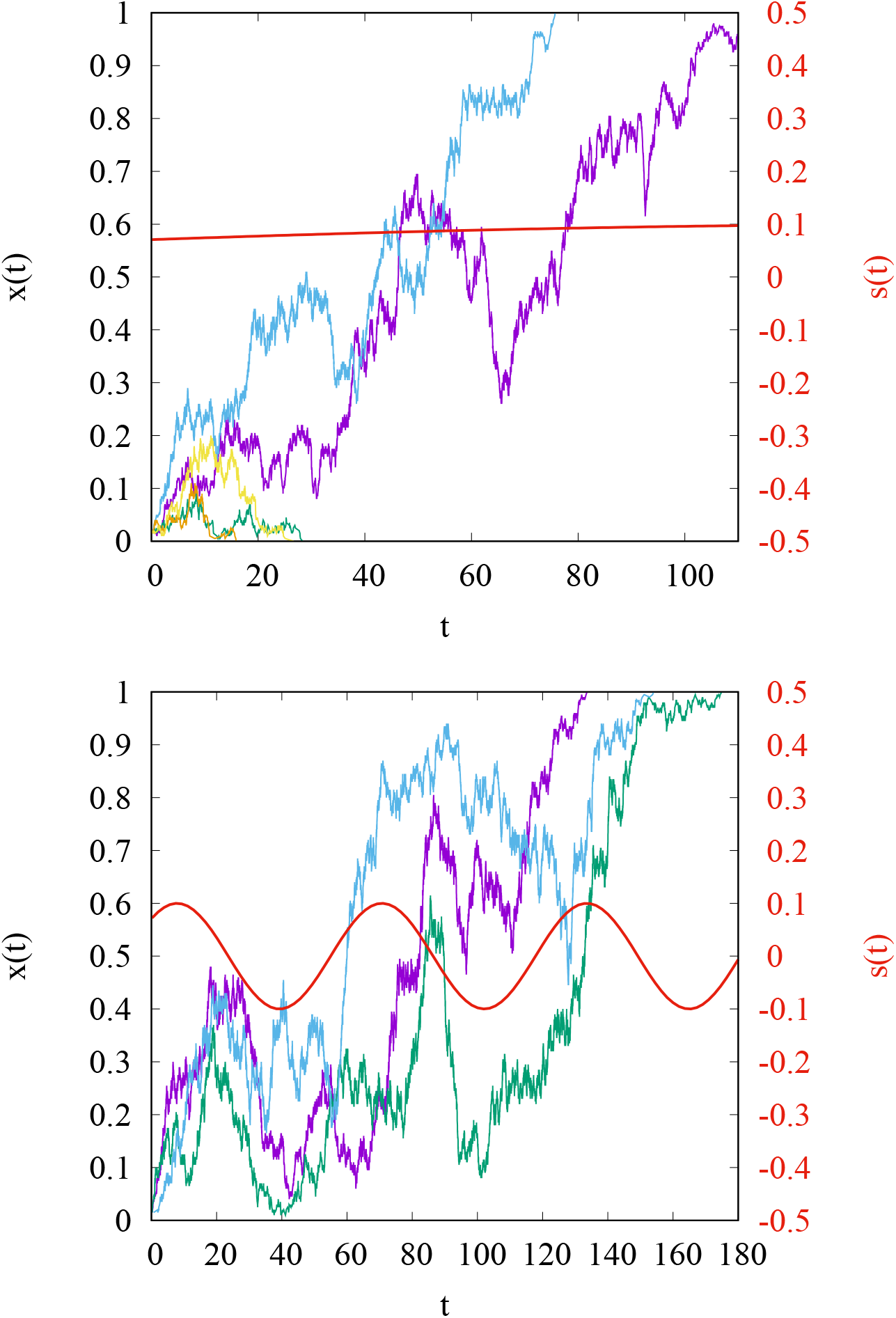
Stochastic trajectories of allele frequency for an on average neutral mutant, starting with a single mutant for *N* = 200, 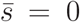, *σ* = 0.1, *h* = 0.5, *θ_a_* = *π*/4 for *ω* = 0.005 (top) and 0.1 (bottom). The successful mutant passes through several cycles of selection before fixing in a rapidly changing environment.

However, in rapidly changing environments (*ω* → ∞), as Fig. 1, Fig. 2 and Fig. 3 demonstrate, the impact of dominance on the fixation probability can be different from that in a static environment. For large cycling frequencies, although the fixation times are much longer than the time period of environmental change (see Fig. 4b), the time of appearance still plays an important role in determining the chance of fixation as the mutant must escape the stochastic loss at short times (in fact, Fig. 4b shows that the effect of random genetic drift is strongest in the first seasonal cycle). Then a mutant that arises in an on average neutral environment while selection is positive but the selection strength is decreasing will soon encounter an environment with negative fitness effects that affect a dominant mutant more adversely than the recessive one leading to a lower chance of fixation of the dominant mutant. In an environment that is beneficial on average, if a mutant arises while selection is negative, it will spend relatively less time in the negative cycle and therefore will effectively behave like a beneficial mutant resulting in the behavior of the fixation probability different from that in the static environment; a similar argument holds when 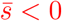.

### 3.2 On average beneficial mutant in a large population

When the mutant is beneficial on average 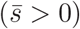 and the population is large enough 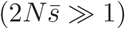, one can use a branching approximation to find the fixation probability of a rare mutant. The basic idea is that a beneficial mutant will get fixed once it is present in finite frequency in the population but it must survive the loss due to random genetic drift when it is initially present in small number compared to the population size. Then the probability that *i* ≪ 2*N* number of *a* alleles are present at time *t* is governed by 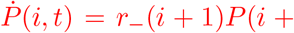 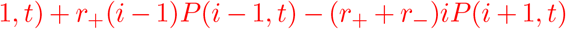 along with boundary condition *P* (*i, t*) = 0, *i* < 0, where *r*_−_(*t*) = lim_*N*→∞_*r_d_/i* and *r*_+_(*t*) = lim_*N*→∞_*r_b_/i* are, respectively, per capita birth rate and death rate of the mutant in a large population. The probability of eventual fixation, *P*_fix_ = 1 − lim_*t*→∞_*P* (0, *t*) is given by (Kendall, 1948)

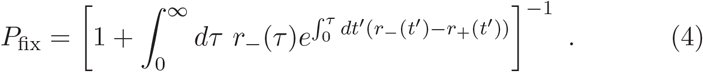

From (1) and (2), we obtain *r*_−_ = 1, *r*_+_ = 1 + *hs*; using these in (4), we find that

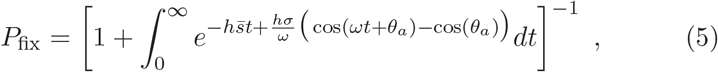

which reduces to (29) of Uecker and Hermisson (2011) for *ξ* = 1 when *h* = 1/2 and *s*(*t*) → 2*s*(*t*) (note that their expression contains a typographical error). Numerical studies of (5) have shown that the fixation probability is, in general, a nonmonotonic function of cycling frequency *ω* and strongly depends on the phase, *θ_a_* = *ωt_a_* (Uecker and Hermisson, 2011; Peischl and Kirkpatrick, 2012).

Equation (5) is analyzed in Appendix A for 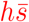, *hσ* ≪ 1, and we find that

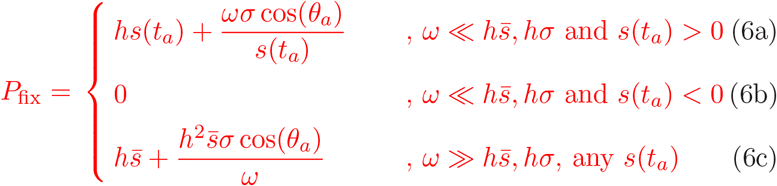

where 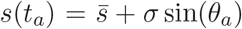. Since the fixation probability of a beneficial mutant with constant selection coefficient *s*_0_ is given by *hs*_0_ (Haldane, 1927), the above expressions show that for slowly changing environments, the fixation probability is determined by the selection coefficient of the mutant at the instant it arose while for rapidly changing environments, it depends on the time-averaged selection coefficient 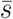 (Gillespie, 1993; Mustonen and LÄssig, 2008).

For 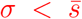, the mutant is beneficial at all times; in this case, the effect of a slowly changing environment is captured by the deviation from *hs*(*t_a_*) in (6a) which changes linearly with the cycling frequency and is *independent* of the dominance parameter. In contrast, for rapidly changing environments, the fixation probability (6c) is sensitive to dominance as the deviation from the asymptotic result 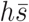 depends on *h* (also, see Fig. S1). The inset of Fig. S1 as also (6a) and (6c) show that the dominant mutant has higher fixation probability than the recessive one at *all* cycling frequencies. In other words, Haldane’s sieve (Haldane, 1927) that favors the establishment of beneficial dominant mutations in static environments continues to operate in changing environments in which the mutant is beneficial at all times. Equations (6a) and (6c) also emphasize the important role of the time of appearance of the mutant. If the beneficial mutant arises while the selection coefficient is increasing (decreasing) with time, the fixation probability at small cycling frequencies increases (decreases) with *ω* and approaches the asymptotic value 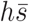 from above (below) at high cycling frequencies.

As explained in Appendix A, on equating the expressions (6a) and (6c), the fixation probability is found to have an extremum at a *resonance frequency*,

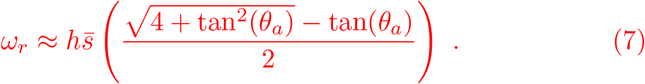

The above expression shows that a minimum or a maximum in the fixation probability occurs when the environment changes at a rate proportional to the average growth rate (Malthusian fitness), 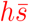 of the population. As already mentioned above, this extremum is a minimum if the mutant appears while selection is decreasing (*π*/2 < *θ_a_* < 3*π*/2) and a maximum otherwise. For the two special values, 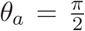 and 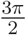, the fixation probability monotonically decreases and increases, respectively, with *ω*.

For 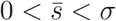 the mutant is not beneficial at all times; in this case, the expression (6a) for small cycling frequencies holds if *s*(*t_a_*) > 0 (see Appendix A). Otherwise as the mutant is initially deleterious and arises in an infinitely large population, the fixation probability is essentially zero. As shown in Fig. 1, the dominant mutant has a higher fixation probability than the recessive one at large cycling frequencies, irrespective of the time at which the mutant appeared (see also (6c)). But at small cycling frequencies, the effect of dominance depends on whether the mutant is beneficial or deleterious when it originated: for positive (negative) *s*(*t_a_*), the dominant (recessive) mutant is favored. For an initially beneficial mutant, as discussed above, the fixation probability changes nonmonotonically with cycling frequency and the resonance frequency is given by (7). But it can exhibit an extremum for initially deleterious mutant also (see Fig. 1 for *θ_a_* = 7*π*/4). Note that the fixation probability curves intersect for different dominance curves which can be estimated for large cycling frequencies as discussed in Appendix A.

To summarize, in an environment that is beneficial on average, the impact of dominance on fixation probability is different in slowly and rapidly varying environments if the mutant is deleterious when it appears in the population.

### 3.3 On average neutral mutant in a finite population

We now calculate the fixation probability of an on average neutral mutant using the backward diffusion equation (3).

#### 3.3.1 Small population

We first consider a small population of size *N* ≪ *σ*^−1^ and analyze the *ω* ≪ *σ* and *ω* ≫ *σ* regimes in Appendix B and Appendix C, respectively. On using (B.8) and (C.2), we find that the fixation probability of allele *a* present in a single copy at time *t_a_* = *θ_a_/ω* is given by

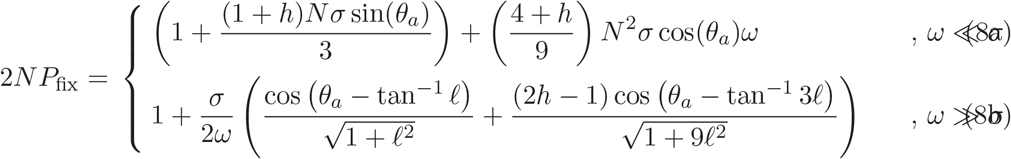

where *ℓ* = (*Nω*)^−1^. The expression (8b) holds for any *ω* ≫ *σ* but for *ω* ≫ *N* ^−1^, it simplifies to

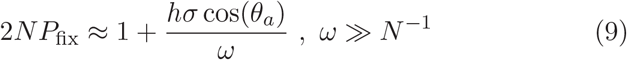

which monotonically approaches the fixation probability of a neutral mutant in a static environment. The above results show that the change in the fixation probability depends weakly on the dominance parameter when the environment changes slowly, but has a strong dependence on *h* in rapidly changing environments. The top panel of Fig. 5 shows that the expressions (8a) and (8b) are in very good agreement with the simulation results, and suggests that the resonance frequency where the fixation probability has an extremum does not depend on the dominance coefficient. As detailed in Appendix C, we find that

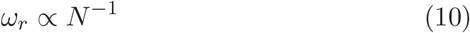

and depends weakly on the dominance coefficient.

**Figure 5:**
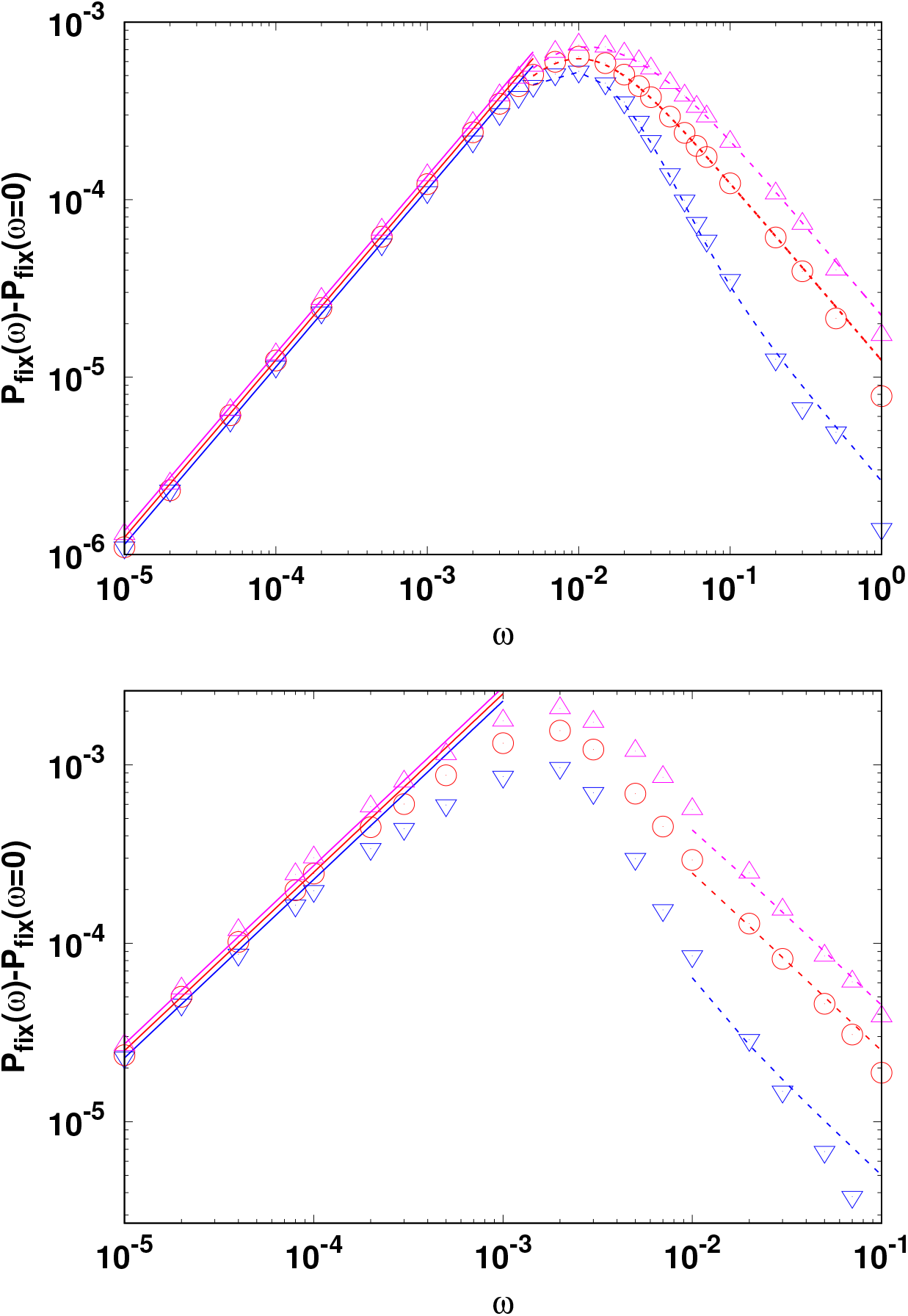
Effect of a changing environment on the fixation probability of a mutant that is neutral on average and arises in a small population (*N* ≪ *σ*^−1^, top) and large population (*N* ≫ *σ*^−1^, bottom). The points show the simulation data obtained by averaging over 10^7^ independent runs. The analytical results obtained using the diffusion theory are shown by lines; the solid lines show the expression (8a) for small cycling frequencies and the dashed lines represent (8b) for large cycling frequencies. Here *N* = 100, *σ* = 0.005 (top) and *N* = 1000, *σ* = 0.01 (bottom), *θ_a_* = 0, and *h* = 0.1(▽), 0.5(◦), 0.9(△). The value *P*_fix_(*ω* = 0) subtracted on the *y*-axis was obtained numerically.

#### 3.3.2 Large population

We now consider large populations with size *N* ≫ *σ*^−1^. For *ω* ≪ *N* ^−1^ ≪ *σ*, as discussed in Appendix B, we find that in slowly varying environments,

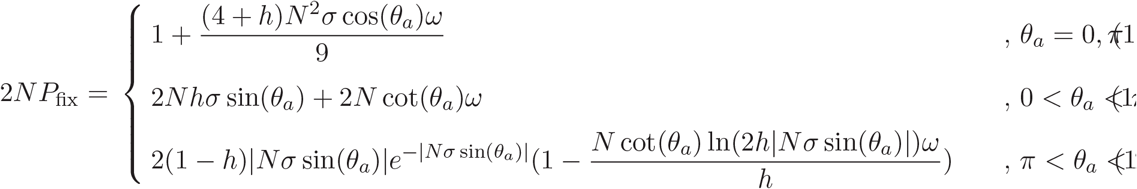

The above equations show that the fixation probability increases linearly with cycling frequency if the mutant arises while selection is increasing but decreases otherwise. For positive *s*(*t_a_*), the magnitude of the slope is independent of *h* and *σ* but varies with these parameters for *s*(*t_a_*) ≤ 0. For large cycling frequencies (*ω* ≫ *N* ^−1^, *σ*), the fixation probability is given by (9) for any *s*(*t_a_*), and approaches the asymptotic neutral behavior from above (below) when 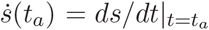 is positive (negative) with the fixation probability increasing (decreasing) with increasing *h*. Thus, as shown in Fig. 2, in an on average neutral environment, the impact of dominance depends on both *s*(*t_a_*) and 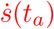. Figure 5 shows a comparison between our analytical and numerical results when the mutant appears at *θ_a_* = 0 (for other values of *θ_a_*, see Fig. S2), and we find a good agreement.

Our perturbation expansions in Appendix B and Appendix C are not valid for intermediate cycling frequencies (*N*^−1^ ≪ *ω* ≪ *σ*). However, our numerical simulations suggest that as for small populations, the resonance frequency scales as *N* ^−1^ here also.

### 3.4 On average deleterious mutant in a finite population

When a mutant is deleterious at all times 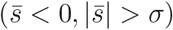, its fixation probability is lower than that of a neutral mutant in both static and time-dependent environments, and the dominant mutant has a lower chance of fixation than the recessive one. But when 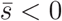 and 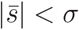, the mutant can be beneficial for some time in a periodically changing environment and its fixation probability can exceed the neutral value depending on the time of appearance. In the left panel of Fig. 3, the selection coefficient *s*(*t_a_*) < 0 and therefore the recessive mutant is favored at small cycling frequencies, while in the right panel of Fig. 3, since *s*(*t_a_*) > 0, the dominant mutant has a higher chance of fixation in slowly changing environments. In either case, at high cycling frequencies, the fixation probability of a recessive mutant is higher than that for a dominant mutant since the time-averaged selection coefficient is negative. Thus, when 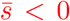, the effect of dominance is different in slowly and rapidly changing environments when *s*(*t_a_*) > 0. We also note that in Fig. 3a, there is a regime where the dominant mutant’s fixation probability exceeds that of the recessive one; however, the difference is quite small and a more detailed investigation is needed to evaluate the importance of this effect.

## 4 Average allele frequency and population fitness

We now turn to the strong mutation regime where 2*Nμ* ≫ 1 and briefly study the dynamics of the allele frequency in changing environments. For 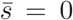, the frequency distribution Φ(*x, t*) of allele *a* under changing selection, mutation, and random genetic drift obeys the following forward Kolmogorov equation (Ewens, 2004),

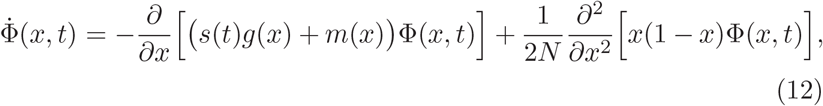

where the mutation term *m*(*x*) = *μ*(1 − 2*x*) and, as before, *s*(*t*) = *σ* sin(*ωt*), *g*(*x*) = *x*(1 − *x*)(*x* + *h*(1 − 2*x*)). The above equation is analyzed in Appendix D for small selection amplitude *σ*, and at large times, the allele frequency distribution Φ(*x, t*) is given by (D.5).

To get an insight into how the allele frequency changes in changing environments, we find the population-averaged allele frequency,

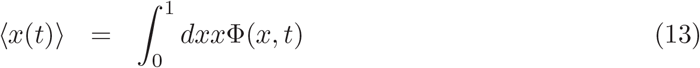

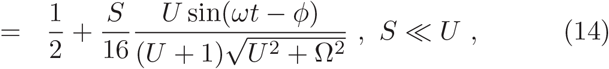

where *U* = 4*Nμ*, *S* = 4*Nσ*, Ω = 2*Nω*, 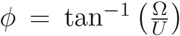. The above equation shows that 〈*x*(*t*)〉 oscillates around one half with the same cycling frequency as *s*(*t*) but a different phase. As depicted in Fig. 6, the phase difference *φ* decreases with increasing mutation rate so that the allele frequency changes almost in-phase with the environment for *U* ≫ Ω but lags behind by a phase *π*/2 for *U* ≪ Ω. The latter behavior for rare mutations is already illustrated in Fig. 4 where the mutant’s allele frequency keeps increasing as long as the selection is positive and decreases when *s*(*t*) becomes negative. In contrast, for *U* ≫ Ω, the population keeps up with the environment as mutations occur faster than the time scale of environmental change.

**Figure 6:**
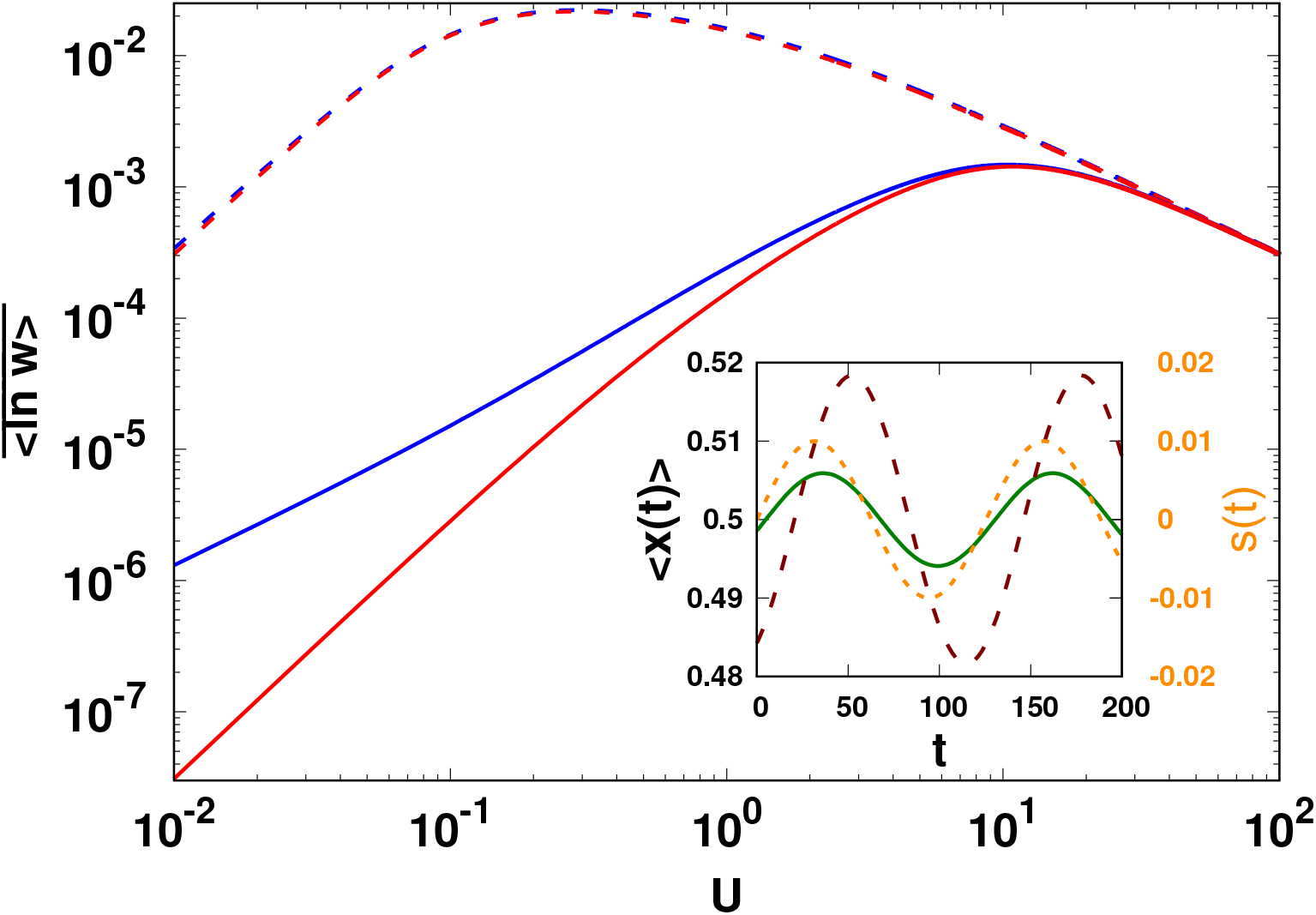
Main: Average log fitness (16) in a periodically changing environment as a function of scaled mutation rate *U* = 4*Nμ* for scaled selection *S* = 4*Nσ* = 20, scaled cycling frequency Ω = 2*Nω* = 0.1 (dashed lines), 10 (solid lines) and dominance parameter *h* = 0.1 (blue), 0.5 (red). The effect of dominance is apparent only for small cycling frequency in the weak mutation regime. Inset: Dynamics of population-averaged allele frequency (14) for *U* = 6 (dashed), 40 (solid). The mutations decrease the phase lag between the average allele frequency and selection (dotted). Note that 〈*x*(*t*)〉 is independent of dominance coefficient. In the inset plots, *N* = 100, *σ* = 0.01, and *ω* = 0.05.

Equation (14) also shows that the allele frequency amplitude remains close to the time-averaged amplitude when the environment changes rapidly. But it is significantly different from one half in slowly changing environments and varies nonmonotonically with the mutation rate with a maximum at the scaled mutation rate *Û* = Ω^2/3^. We also find that although the distribution Φ(*x, t*) depends on the dominance coefficient (see Appendix D), the average allele frequency (for small *σ*) is independent of *h*.

The above described behavior of allele frequency has implications for the average fitness of the population. When the mutant allele is present in frequency *x* at time *t*, the population fitness *w*(*x, t*) = (1+*s*)*x*^2^ +2*x*(1−*x*)(1+*hs*)+(1−*x*)^2^ = 1+*σ* sin(*ωt*)*f* (*x*) where *f* (*x*) = (1 − 2*h*)*x*^2^ + 2*hx* (see MODEL section). The population-averaged log fitness 〈ln *w*〉 oscillates around a constant which is obtained on averaging over a period of the oscillation and is given by

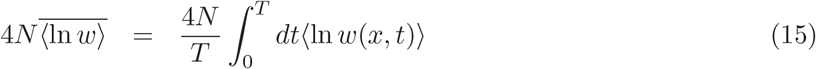

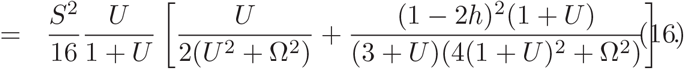

We thus find that although the selection is zero on average, the population fitness is nonzero.

Equation (16) shows that for a given mutation rate, the average log fitness decreases towards zero with increasing cycling frequency; this behavior is expected as the time-averaged environment governs the dynamics in a rapidly changing environment. However, it is a nonmonotonic function of the scaled mutation rate *U*: for 1 ≪ *U* ≪ Ω, the average log fitness is close to zero because the phase difference *φ* between the allele frequency and selection is large (see (14)). But for high mutation rate (*U* ≫ Ω), although the phase lag is small, the allele frequency does not deviate substantially from one half resulting in low fitness. From (16), we find that the average fitness has a peak at an optimal mutation rate,

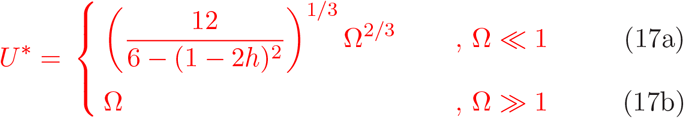

which increases with the rate of environmental change. Finally, equation (16) also shows that the fitness is a symmetric function of dominance coefficient but the *h*-dependence is quite weak (see Fig. 6), and is apparent only at small mutation rates and for fast environmental changes; we thus conclude that dominance does not have an appreciable effect when recurrent mutations occur.

## 5 Data availability

The authors state that all data necessary for confirming the conclusions presented in the article are represented fully within the article.

## 6 Discussion

In this article, we studied the evolutionary dynamics of a finite, diploid population in a varying environment for both weak and strong mutations. The fixation probability of a mutant has been studied in infinitely large populations when selection changes gradually in both magnitude and direction (Uecker and Hermisson, 2011; Peischl and Kirkpatrick, 2012) and in finite populations that are subjected to abrupt changes in the direction of selection (Takahata *et al.*, 1975; Mustonen and LÄssig, 2008; CvijoviĆ *et al.*, 2015; Dean *et al.*, 2017). Here we modeled a situation in which the mutant allele is beneficial during a part of the seasonal cycle and deleterious in another, and hence its selection coefficient *s*(*t*) varies periodically with time. In contrast to previous work, here we focused on the impact of dominance on evolutionary dynamics in changing environments, and obtained simple analytical expressions for the fixation probability in both infinite and finite populations.

### Rate of environmental change and time of appearance

In a slowly changing environment, the fixation probability of a mutant is expected to depend on the time it appears in the population. But it is perhaps not obvious if the dependence on the initial condition remains in fast changing environments as the fixation probability in an infinitely fast changing environment is given by the corresponding result in a static environment with the time-averaged selection coefficient 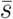. However, as shown in Fig. 4, a mutant must escape the stochastic loss at short times in order to fix in the population, and therefore the eventual fate of the rare mutant depends on its time of appearance at any finite rate of environmental change.

Using branching process and diffusion theory (Uecker and Hermisson, 2011; Waxman, 2011), here we have obtained simple expressions for the fixation probability when the frequency of the environmental change is smaller or larger than the resonance frequency *ω_r_* of the population which is given by the average growth rate of the population when the time-averaged selection strength 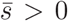 and inverse population size for 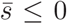. The fixation probability exhibits an extremum when the environment changes at a rate equal to the resonance frequency; whether this extremum is a minimum or a maximum also depends on the time of appearance, *t_a_* of the mutant.

As an illustration of the above discussion, for arbitrary 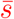, consider a beneficial mutant that arises when its selection strength is decreasing with time. In a slowly changing environment, as 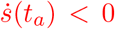, its fixation probability is expected to be smaller than that in the static environment; for the same reason, it approaches the corresponding result in the time-averaged environment from below thus resulting in a minimum at the resonance frequency. It then follows that in a periodically varying environment with zero or negative time-averaged selection coefficient, if the environment changes at a rate faster than the resonance frequency, a mutant that is beneficial in a static environment will have a fixation probability lower than (2*N*)^−1^. Similarly, a deleterious mutant that arises while selection strength is increasing can have enhanced chance of fixation compared to (2*N*)^−1^ in changing environments that are neutral or beneficial on average.

The above discussion assumes that the mutations are rare. When recurrent mutations occur, we find that in the on average neutral environment, the population can gain fitness which is, however, appreciable when the mutation rate is as high as the rate of environmental change.

### Role of dominance in slowly changing environments

In a static environment, a dominant beneficial mutant enjoys a higher chance of fixation than a recessive one because the (marginal) fitness of the mutant allele (relative to the wild type allele) is higher in the former case (Haldane, 1927). This behavior is reversed for deleterious mutants where the fixation of recessives is favored (Kimura, 1957). In a slowly changing environment (*ω* ≪ *ω_r_*), if the mutant starts out as a beneficial mutant (that is, its selection coefficient at time of appearance *s*(*t_a_*) > 0), Haldane’s sieve operates. Similarly, if the mutant is deleterious to begin with, its chances of fixation are reduced if it is dominant. This result is attested by Figs. 1 and S1 for 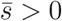, Fig. 2 for 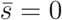 and Fig. 3 for 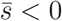.

Equations (6a) and (11b) for the fixation probability of a mutant in an environment that is, respectively, beneficial and neutral on average suggest that the change in fixation probability due to a slow change in the environment is simply equal to the change in the mutant’s initial fitness relative to its initial fitness, that is, 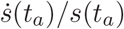 which is *independent* of the dominance coefficient. This can be argued as follows: when the selection coefficient changes very slowly (*ω* → 0), it is reasonable to assume that the fixation probability has the same functional form as that in the static environment. Then for an initially beneficial mutant, 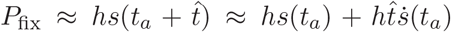 where 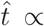 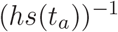 is the time at which the mutant escapes stochastic loss, as estimated from a deterministic argument.

### Role of dominance in fast changing environments

As already mentioned above, an initially beneficial mutant arising when the selection is declining can behave effectively as a deleterious mutant in a rapidly changing environment which is neutral or deleterious on average. This has the immediate consequence that the dominant mutant is less likely to fix than the recessive one as supported by Fig. 2 for 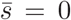 and Fig. 3 for 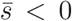. This result can be relevant to understanding adaptation in environments that change fast and for a short period of time. In Sec. S4, we construct such examples and find that the fixation probability of an initially beneficial mutant in transiently changing environments exhibits the same dependence on dominance coefficient as discussed above for periodically changing environments.

Equations (6c) and (9) show that in fast changing environments, the fixation probability is proportional to 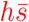 and (2*N*)^−1^, respectively, which are the results for the fixation probability of a single mutant in an infinitely fast changing environment. If genetic drift is ignored, the dynamics of the mutant frequency are described by 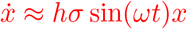 so that at large times, the average number of mutants, *n* = 1+*hσ* cos(*θ_a_*)*/ω* for large *ω* and therefore, the fixation probability in rapidly changing environments can be interpreted as simply that of *n* mutants in infinitely fast changing environments (also, see Sec. S4).

### Limitations and open questions

Our analytical results that provide an understanding of conditions under which the effect of dominance on the fixation probability is different in static and changing environments are applicable to cycling frequencies that are much smaller or larger than the resonance frequency. As our analysis is not valid for intermediate cycling frequencies, we can not rule out if such differences occur at cycling frequencies around resonance frequency (see, Fig. 1 for *θ_a_* = 7*π*/8 and Fig. 3a).

Here we have mainly focused on the fixation probability and did not discuss how substitution rate and adaptation rate behave in changing environments. However, our preliminary simulations show that the substitution rate varies nonmonotonically with cycling frequency (also see Mustonen and LÄssig (2007)). A detailed understanding of these quantities requires the knowledge of fixation time which shows interesting dependence on dominance coefficient in static environments (Mafessoni and Lachmann, 2015); extending such results to temporally varying environments is desirable and will be discussed elsewhere.

When adaptation occurs due to standing genetic variation, the fixation probability of a beneficial mutant is known to be independent of dominance in static environments (Orr and Betancourt, 2000); here, we have studied the fixation probability of a *de novo* mutation and a detailed understanding of how standing variation affects the results obtained here is a problem for the future.

## Acknowledgments

We thank two anonymous reviewers for many constructive comments that helped us to improve the manuscript.

## Appendix A Branching process approximation

The fixation probability of a mutant that arises in a large wild type population and is beneficial on average is given by (5). For *ω* → ∞, the fixation probability is given by 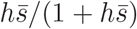 while for *ω* = 0, it is equal to *hs*(*t_a_*)/(1 + *hs*(*t_a_*)) for *s*(*t_a_*) > 0, and zero otherwise.

Away from these extreme limits, the integral appearing in (5) can be analyzed for small 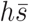, *hσ* as follows. For small cycling frequencies 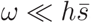, *hσ*, by first expanding the integrand in powers of *ω* and then carrying out the resulting integrals, we obtain

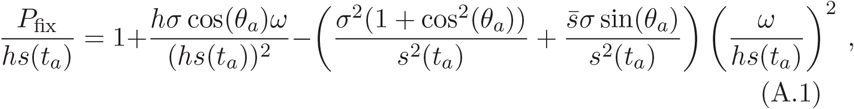

provided *s*(*t_a_*) > 0, and zero otherwise. Similarly, for large cycling frequencies, the fixation probability can be found by first expanding the integrand in powers of *hσ/ω*; for 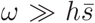, *hσ*, this finally results in

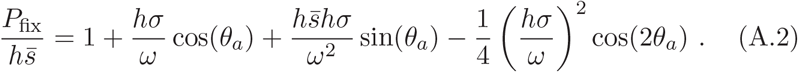

Note that the first order corrections in (A.1) and (A.2) vanish for *θ_a_* = *π*/2 and 3*π*/2.

As discussed in the main text, an extremum in the fixation probability occurs at the resonance frequency *ω_r_*. Ignoring terms of order *ω*^2^ and 1*/ω*^2^ in (A.1) and (A.2), respectively, and equating the resulting expressions, we arrive at a quadratic equation for *ω_r_* whose positive root is given by (7). Furthermore, in Fig. 1, for *π*/2 < *θ_a_* < 3*π*/2, the fixation probability curves for dominance coefficient *h* and *h*′ coincide at a cycling frequency higher than *ω_r_* which can be estimated using (A.2), and found to be −(*h* + *h*′)*σ* cos(*θ_a_*).

## Appendix B Diffusion approximation for small cycling frequencies

Here we study (3) for 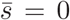 and small cycling frequencies within a perturbation theory by writing *P*_fix_ = *P*_0_ + *NωP*_1_. It is useful to rewrite (3) as

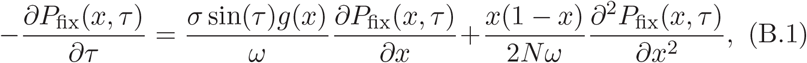

where *τ* = *ωt* + *θ_a_, t* ≥ 0.

In a static environment, if the mutant arises at time *t_a_* = *θ_a_/ω* and has fraction *x* in the population, its fixation probability is given by 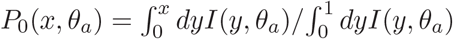, where

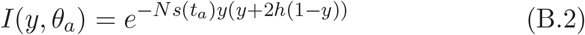

and *s*(*t_a_*) = *σ* sin(*θ_a_*) (Kimura, 1957). For a strongly beneficial mutation (*Ns*(*t_a_*) ≫ 1), the fixation probability *P*_0_ increases with dominance coefficient and given by *hs*(*t_a_*), while for a deleterious mutation, it decreases with *h*. The chance of fixation also decreases with population size for *h* ≤ 1/2 but the variation with *N* is non-monotonic for *h* > 1/2.

The effect of a slowly changing environment on the fixation probability is captured by *P*_1_ that, by virtue of (B.1), obeys the following ordinary differential equation,

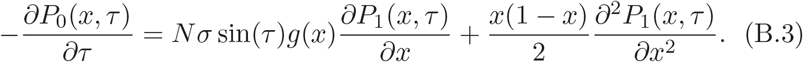

Equation (B.3) subject to boundary conditions *P*_1_(0, *τ*) = *P*_1_(1, *τ*) = 0 has the solution

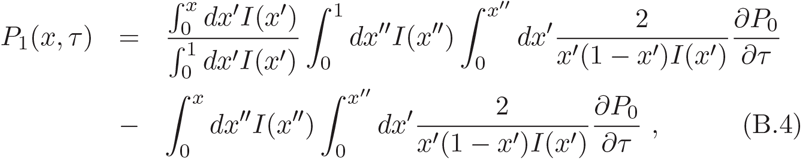

which, for small initial frequency (*τ* = *θ_a_, x* → 0) can be approximated by

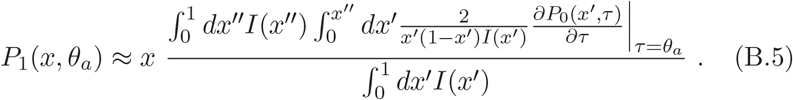

The following cases need to be considered separately:

i. −1 ≪ *Ns*(*t_a_*) ≪ 1: For small *Nσ* and arbitrary *θ_a_*, we first expand *I*(*x, τ*) to linear order in *Nσ* and carry out the integrals in the expression for *P*_0_ given above to obtain

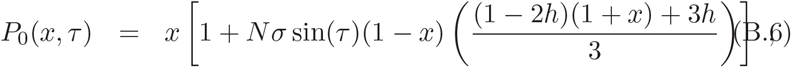

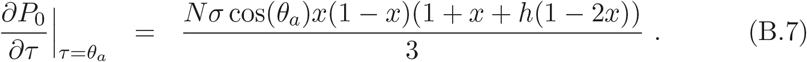 Using these approximations in (B.5), to leading order in *Nσ*, we get

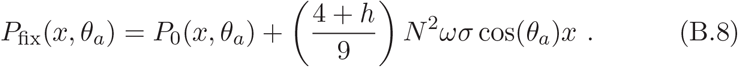 For *Ns*(*t_a_*) = 0 (that is, *θ_a_* = 0 or *π*, arbitrary *Nσ*), it can be easily seen that the function *I*(*x, θ_a_*) = 1 and the derivative 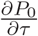 is given by (B.7) thus leading to (B.8).
ii. *Ns*(*t_a_*) ≫ 1: For large |*Ns*(*t_a_*)|, using the asymptotic expansion of the error function erf(*x*) (Abramowitz and Stegun, 1964), the fixation probability *P*_0_(*x, τ*) can be approximated as

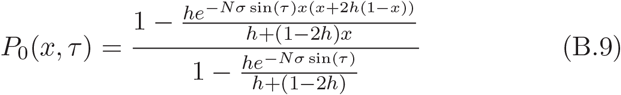

(more precisely, the above expression holds 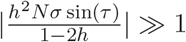). For large, positive *Ns*(*t_a_*), the denominator in (B.9) can be approximated by one leading to

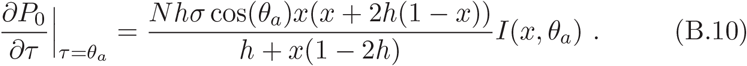 Using the above expression in (B.5) and performing the integrals for *Ns*(*t_a_*) ≫ 1, we finally obtain the following simple result,

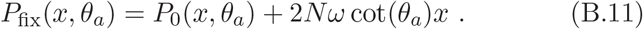
iii. *Ns*(*t_a_*) ≪ −1: Taking the derivative of *P*_0_ in (B.9) with respect to *τ* and keeping factors proportional to *e*^−*Nσ*^ ^sin(*τ*)^ only, we obtain

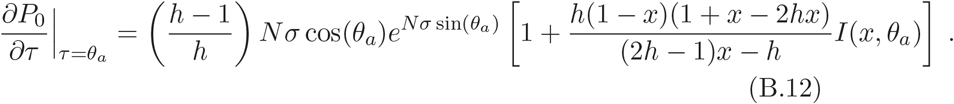

Noting that the dominant contribution to the inner integral in the numerator of (B.5) comes from *x*′ → 0, we finally get

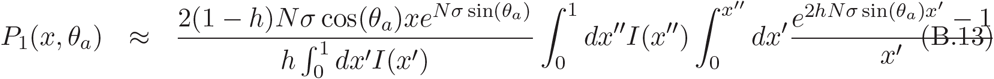

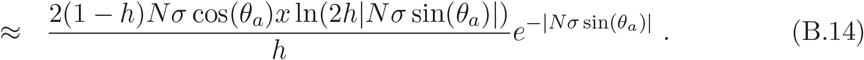

## Appendix C Diffusion approximation for large cycling frequencies

Here we calculate the fixation probability of a neutral mutant 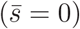 when the cycling frequency is larger than the amplitude of selection (*ω > σ*). On writing 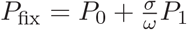 in (B.1) and collecting terms to zeroth and first order in *σ/ω*, we find that *P*_0_ = *x*, as expected. The correction *P*_1_ obeys an inhomogeneous partial differential equation,

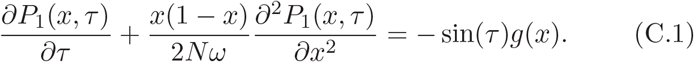

with boundary conditions *P*_1_(0, *τ*) = *P*_1_(1, *τ*) = 0.

The homogeneous equation can be solved using standard eigenfunction expansion method (Kimura, 1955; Ewens, 2004), and we find that 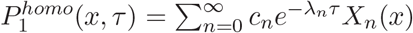 with the eigenvalue 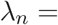 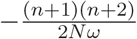 and eigenfunction 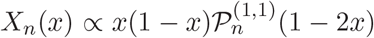 where 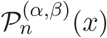 is the Jacobi polynomial (Abramowitz and Stegun, 1964). However, as this homogeneous solution is not periodic in *τ*, it does not contribute to the full solution. But since the eigenfunctions *X_n_*(*x*) form a complete set of basis, we can write *P*_1_(*x, τ*) = Σ_*n*=0_ *a_n_X_n_*(*x*) and *g*(*x*) = Σ_*n*=0_ *b_n_X_n_*(*x*) where *b_n_* are obtained using the orthogonality property of *X_n_*(*x*). Using these in (C.1), we obtain

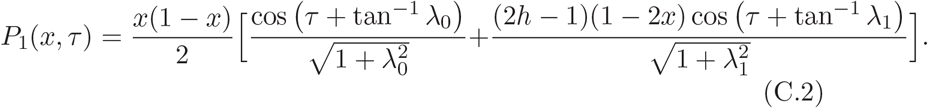

When a single mutant with frequency *x* = (2*N*)^−1^ appears at time *t_a_*, the above expression reduces to (8b) in the main text and can be used to find the resonance frequency at which the probability of fixation has an extremum. For *h* = 1/2, we find that

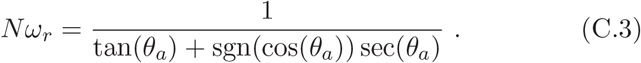

For arbitrary *h*, we are unable to find a simple closed expression for *ω_r_* as it is a solution of a 6th order algebraic equation. But a numerical study of this equation shows that *ω_r_* depends weakly on dominance coefficient. For *θ_a_* = 0, we find *Nω_r_* ≈ 0.88, 1.0, 1.14 for *h* = 0.1, 0.5, 0.9, respectively; the corresponding values for *θ_a_* = 3*π*/4 are given by 1.73, 2.41, 3.84.

## Appendix D Allele frequency distribution for strong mutation

The forward time dynamics of the population under mutation and selection are described by (12) for the allele frequency distribution Φ(*x, t*). The distribution Φ_0_ for the population subject to mutation and random genetic drift is given by 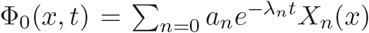 with eigenvalues 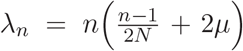 and eigenfunctions 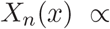 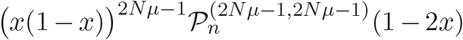 where, 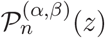 is the Jacobi polynomial (Crow and Kimura, 1956). These eigenfunctions are orthogonal with respect to the weight function *ρ*(*x*) = [*x*(1 − *x*)]^1−2*Nμ*^.

For weak selection (*σ < μ*), we can expand Φ(*x, t*) as a power series in *σ/μ* to write Φ = Φ_0_ + (*σ/μ*)Φ_1_. Using this in (12), we find that Φ_1_ obeys the following differential equation,

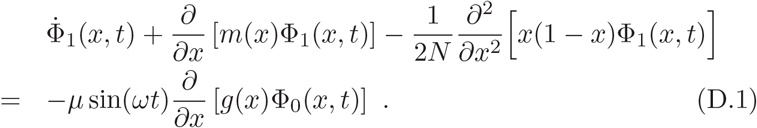

To find the distribution Φ_1_(*x, t*), we expand it and the RHS of above equation as a linear combination of *X_n_*(*x*). Writing 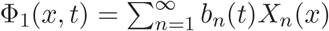 in (D.1), we find that at large times,

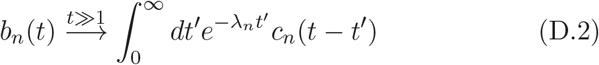

where

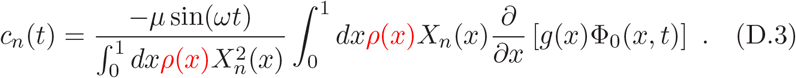

As 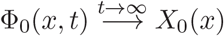, from (D.2), we obtain

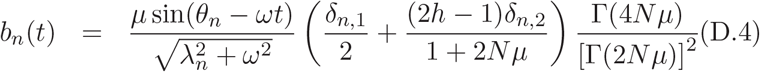

where *θ_n_* = tan^−1^(*ω/λ_n_*). We thus have

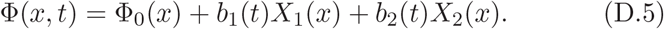

## Appendix S1 Discrete and continuous time models for diploids

Consider a randomly mating population of *N* diploids with discrete generations in which the selection acts on the viability of an individual. Then the life cycle consists of a juvenile phase during which mutation, selection and random genetic drift act, and determine the chance of survival of an individual to the adult phase in which reproduction occurs. For a large population, immediately after random mating, the frequency of the genotypes can be approximated by the corresponding Hardy-Weinberg proportions as the deviations from it are of order 1*/N* (Nagylaki, 1992). The dynamics in the juvenile phase can be modeled by a discrete time Wright-Fisher process (Ewens, 2004).

In contrast to the discrete time model described above, in the main text, we worked with a continuous time model as the environment is assumed to change gradually. In overlapping generations, a model that incorporates details of the life cycle is necessarily complex as the age-structure of the population must be carefully taken into account. Moreover, besides large *N*, additional assumptions, viz., small selection coefficient, *s* and mutation probability, *μ* are required for Hardy-Weinberg equilibrium to hold in such continuous time models (Nagylaki, 1992).

However, in the diffusion approximation where *s* → 0, *μ* → 0, *N* → ∞ with finite 2*Ns,* 2*Nμ* in the continuous time model and 4*Ns,* 4*Nμ* in the discrete time model, we obtain essentially the same Kolmogorov equations for both models (Ewens, 2004). In the main text, we studied the dynamics of adaptation in the framework of diffusion theory when the mutant is on average neutral as the 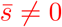 case does not seem to be analytically tractable.

However, for positive 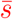, it is possible to make analytical progress by modeling the fixation process as a branching process (Desai and Fisher, 2007; Uecker and Hermisson, 2011). The branching approximation applies as long as the mutants are rare, that is, a finite number of mutants are present in an infinitely large population. In this approximation, the transition rate matrix for the discrete time model discussed above is not a continuant matrix (Ewens, 2004). However, as the number of mutants is small, it is reasonable to assume that the transition rates are significantly different from zero only when the number of mutant allele changes by one in a generation; in other words, we arrive at a birth-death model. The above discussion thus provides a justification for the birth-death model for diploids and in the main text, we have used it for all parameter regimes.

## Appendix S2 Fixation probability of a mutant that is beneficial at all times

**Figure S1:**
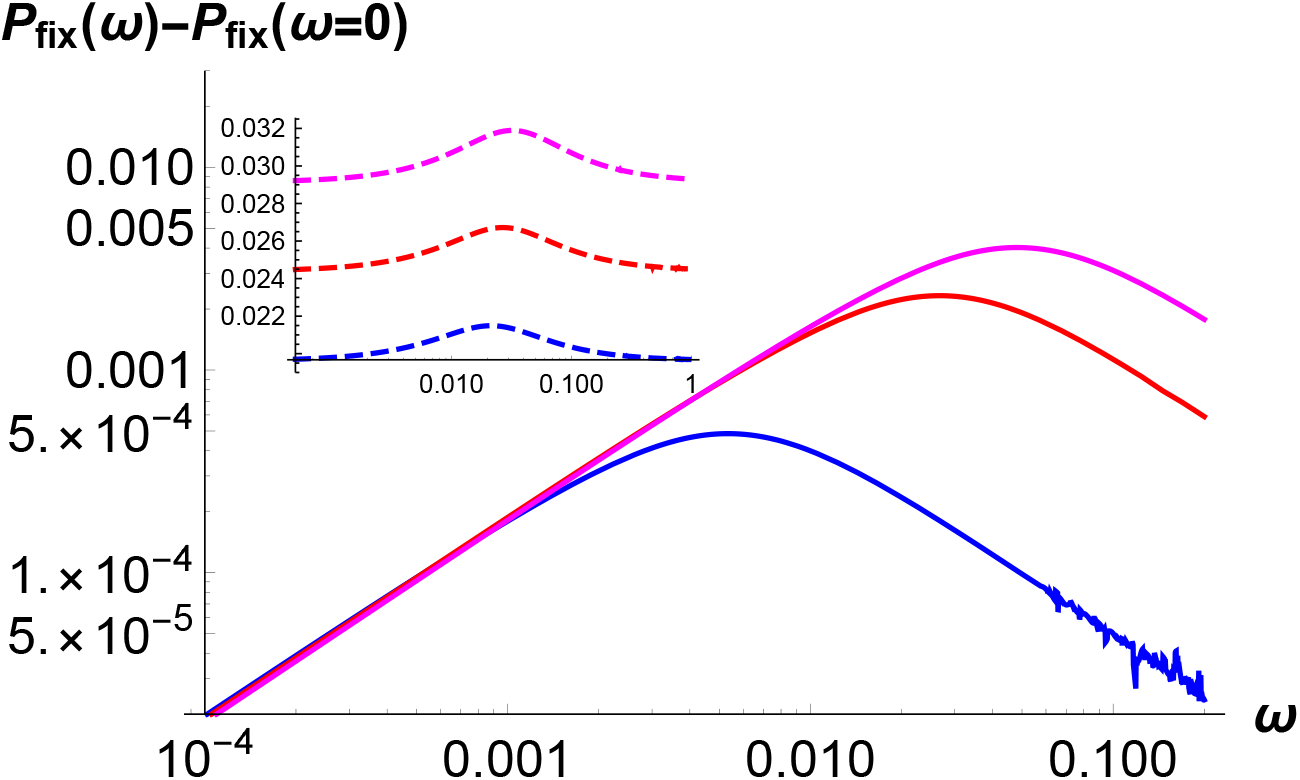
The inset shows the fixation probability *P*_fix_(*ω*) given by (5) for a mutant that is beneficial on average for dominance coefficient *h* = 0.4, 0.5, 0.6 (bottom to top). In the main figure, the effect of a changing environment is shown by subtracting the fixation probability *P*_fix_(*ω* = 0) = *hs*(*t_a_*)/[1 + *hs*(*t_a_*)] with 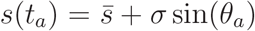 for *h* = 0.1, 0.5, 0.9 (bottom to top). In both plots, 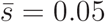, 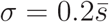, *θ_a_* = 0.

## Appendix S3 Fixation probability of an on average neutral mutant

**Figure S2:**
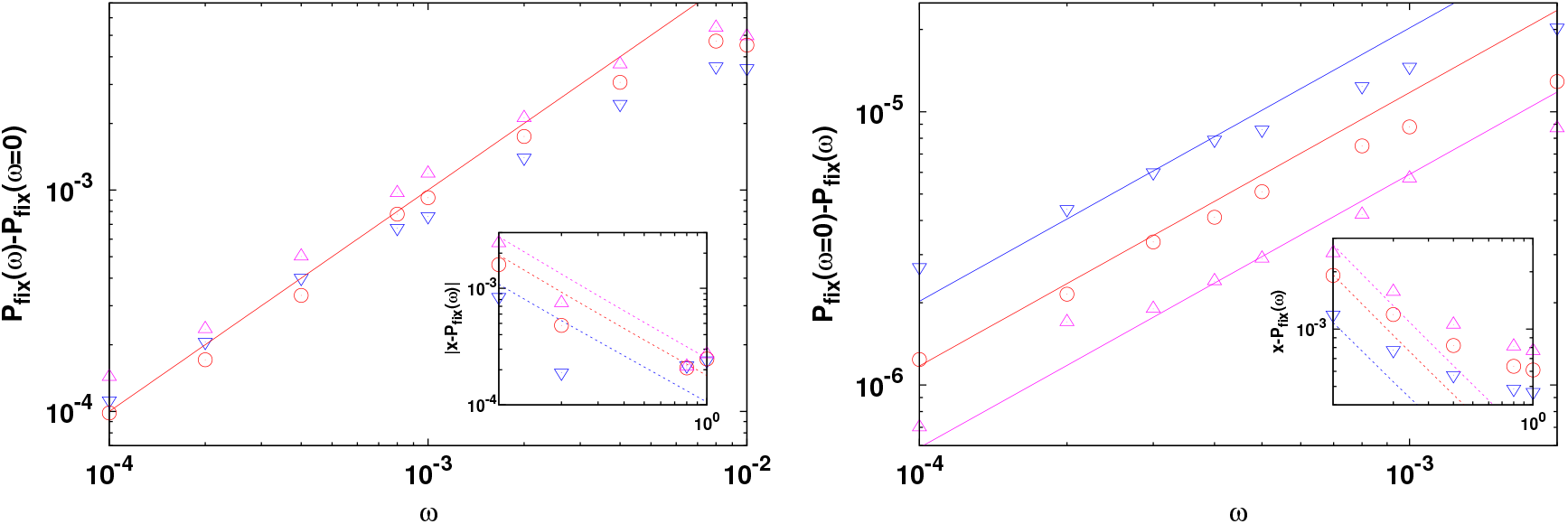
Fixation probability of a mutant that is neutral on average for *Nσ* ≫ 1 when the mutant appeared at *θ_a_* = *π*/4 (left panel) and 5*π*/4 (right panel). The solid line shows the expression (11b) (left panel) and (11c) (right panel) for small cycling frequencies and the dashed lines represent (8b) for large cycling frequencies. Here *N* = 100, *σ* = 0.1 and *h* = 0.3(▽), 0.5(∘), 0.7(△). The numerically obtained value *P*_fix_(*ω* = 0) = 2.53 × 10^−2^, 3.40 × 10^−2^, 4.38 × 10^−2^ for *h* = 0.3, 0.5, 0.7, respectively, for the left panel. For the right panel, *P*_fix_(*ω* = 0) = 3.55 × 10^−5^, 2.58 × 10^−5^, 1.98 × 10^−5^ for *h* = 0.3, 0.5, 0.7, respectively. The simulation results are averaged over 10^7^ runs.

## Appendix S4 Fixation probability of a mutant in transiently varying selection

**Figure S3:**
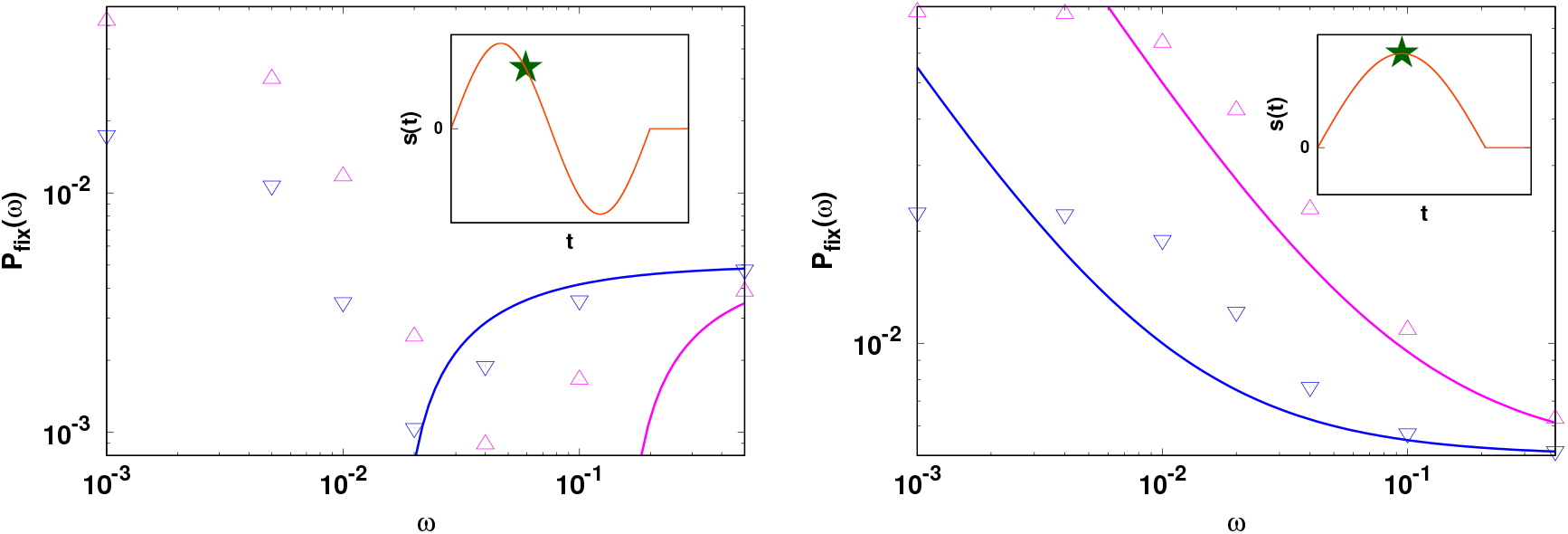
Fixation probability of a mutant in transiently varying environments defined by (S4.1) and (S4.2) and illustrated in the inset. The lines show the expression (S4.4) and the points show the simulation data obtained by averaging over 10^7^ independent runs for *θ_a_* = 3*π*/4, *T_e_* = 2*π/ω* (left panel) and *θ_a_* = *π*/2, *T_e_* = *π/ω* (right panel). In all the plots, *N* = 100, *σ* = 0.1 and *h* = 0.1(▽), 0.9(△).

Here we consider a situation in which the time-averaged selection coefficient is nonzero and changes over a finite time *T_e_*,

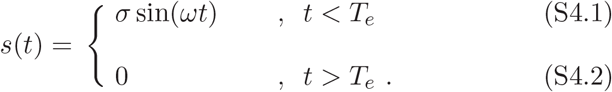

Waxman (2011) has shown that in such a case, the fixation probability is simply given by the mean allele frequency at the end of selection. Here we estimate this allele frequency using the deterministic evolution equation,

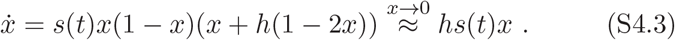

For large *ω*, starting from a single mutant, the number of mutants at time *T_e_* is then given by

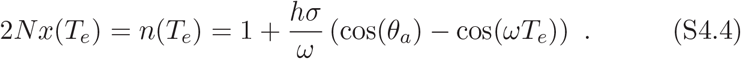

For large cycling frequencies, this prediction matches qualitatively with the numerical results in Fig. S3, and therefore captures the effect of dominance when the environment changes fast over a short interval of time.

